# Automating the Curation of DNA Barcode Databases for Vascular Plants

**DOI:** 10.1101/2025.04.22.650100

**Authors:** Andreas Kolter, Paul Hebert

**Affiliations:** Centre for Biodiversity Genomics, University of Guelph, Guelph, ON N1G 2W1, Canada

## Abstract

Comprehensive, curated, and current DNA barcode reference databases are essential for both the identification of single specimens and for the interpretation of metabarcoding data. In the case of plants, nuclear (ITS) and plastid (rbcL, matK) markers are commonly utilized in union. Because the plastid regions are segments of protein-coding genes, their alignment and analysis are usually straightforward. By contrast, the assembly and validation of records for ITS is considerably more difficult for two reasons – the prevalence of indels and the presence of intraindividual variation. This complexity has provoked the development of several workflows to support the curation of reference databases for the internal transcribed spacer (ITS) region for plant barcoding. However, the pipelines used to create these databases lack functionalities which are essential to ensure a solid post-analytical validation. This paper presents a new workflow to address these shortcomings, with the goal of enhancing the reliability and accuracy of plant barcoding studies. We furthermore demonstrate that clustering of reference databases results in a substantial drop in the fraction of queries that gain a correct species-level assignment. By contrast, setting an acceptance threshold for identifications, based on the distance between query and match provides a meaningful reduction of error rates in incomplete reference databases.

## Introduction

DNA barcoding offers an alternative to morphological identification methods (Kress, 2017). Instead of relying on morphologically diagnostic traits, plant tissue from any developmental stage can be used to infer a taxonomic identity. Plant metabarcoding builds on this concept and expands the methodology to samples that derive from multiple species. This latter approach facilitates dietary analysis and ecological interactions using matrices such as stomach contents or fecal matter (Hollingsworth et al., 2011; Bruni et al., 2015; Kartzinel et al., 2015). Other important applications are reconstructing plant species assemblages from ancient sediment, identifying food adulteration, validating herbal medicine, and tracking airborne pollen loads (Bruno et al., 2019; Urumarudappa et al., 2020; Huang et al., 2021; Krinitsina et al., 2023).

It is crucial to have the ability to compare the results from any barcoding study with a comprehensive, curated and current DNA barcode reference database. Ideally, the reference database should include multiple sequences of every plant species obtained from well-identified, vouchered plant specimens (Kolter and Gemeinholzer, 2021a). At present, reference databases are far from this ideal as more than 80% of flowering plant species either lack coverage entirely or have few records. Among the molecular markers used for plant barcoding, the internal transcribed spacer (ITS) region within the nuclear ribosomal DNA array (nrDNA), has factually become a core plant barcode (China Plant BOL Group et al., 2011). GenBank (Sayers et al., 2019) provides the most comprehensive, easily accessed source of plant ITS sequences. However, these records lack consistent annotation, are not screened for pseudogenes, and misidentifications are prevalent. RefSeq, a curated database also hosted by the National Center for Biotechnology Information (NCBI), partially addresses these deficits, but it only encompasses fungal ITS (O’Leary et al., 2016). Until now, only Canada (Kuzmina et al., 2017) and the UK (Jones et al., 2021) have published near-complete ITS reference libraries for use in a national context. In summary, a comprehensive, reliable DNA barcoding reference database for plants currently requires downloading and filtering GenBank data.

Multiple pipelines to create DNA barcode reference databases already exist, each with unique attributes. Four general workflows that can be applied to plants and other markers are: 1) BCdatabaser, which focuses on user-friendly execution and automatic uploads to Zenodo (Keller et al., 2020); 2) rCRUX which uses an iterative BLAST approach to identify unannotated marker sequences (Curd et al., 2023); 3) CRABS combines in-silico PCR with an alignment to identify marker sequences for inclusion in the reference database (Jeunen et al., 2023); and 4) MetaCurator uses hidden Markov models to extract the desired marker region from a set of sequences (Richardson et al., 2020).

However, in general, plant-specific curatorial steps are necessary to deal with ITS pseudogenes and a lack of ITS primer specificity for plants, which can result in off-target amplification (Kolter and Gemeinholzer, 2021b; Zhang et al., 2022). Although exposure to these complexities can be reduced by wet lab protocols, such as the use of plant-specific primers or adjustments in PCR conditions to avoid pseudogene amplification (Buckler et al., 1997; Cheng et al., 2016), these methods are rarely employed. Plant-specific curatorial steps are implemented in the following three protocols: 1) The static ITS Database V uses secondary structure validation to verify ITS2 sequences (Ankenbrand et al., 2015); 2) PLANits uses the software ITSx to annotate ITS marker sequences (Bengtsson-Palme et al., 2013; Banchi et al., 2020); 3) A database curation script by Quaresma et al. (2023) uses a dynamically executed workflow implementing plant-specific filter steps.

The present script was developed in response to the fact that current solutions lack functionalities essential to post-analytical validation and fail to retain sequence metadata. The novel combination of features introduced here are: (1) automated taxonomic curation, (2) GenBank metadata retention, (3) occurrence information, and (4) the automatic addition of fungal sequences as a sink for contaminated query sequences. We further investigated the impact of database clustering and the exclusion of database matches based on distance thresholds on the success in securing a species-level identification.

## Results

The ITS reference database for plants generated by the present script on 20 Nov 2023 (10.5281/zenodo.10257823) encompasses 271,418 records, each indexed by its GenBank accession number (Table 1). Every record includes 40 columns of metadata and DNA sequences (Supplementary Table 1). Over 80% of ITS2 and 90% of ITS1 sequences were derived from the full ITS (226,407) sequence annotations (Table 1). In addition, the database includes sequences for three additional nrDNA regions (110,039 partial 18S, 234,477 full 5.8S, 86,003 partial 26S), which can be used for primer design or custom data validation. Approximately 70% of all records have voucher information (defined by the sequence owner), extracted from GenBank, linking the sequence to a physical sample (10.5281/zenodo.10257823). Overall, 86% of species possessed at least one data record with voucher information. The database lists 19,945 unique author names and 10,907 unique publications. Almost all records (98%) have at least one GBIF occurrence (summarized as country and count). Comparison of the species names associated with the accessions downloaded from GenBank indicated that approx. 12% (12,861 / 107,710) are regarded as synonyms by GBIF. NCBI taxonomy queries to the GBIF backbone taxonomy yielded a high match rate, with an average confidence score of >99.5% for returned matches.

**Table 1:** Database summary counts by filtering steps.

| step | subset | sequences | species | genera | families | order |
| --- | --- | --- | --- | --- | --- | --- |
| GenBank download | all records | 437414 | 115337 | 11501 | 455 | 78 |
| species-level ID & ambiguous nucleotide filter | all records | 409034 | 112545 | 11339 | 449 | 77 |
| ITSx | ITS1 | 328493 | 104283 | 10747 | 417 | 75 |
|  | ITS2 | 358985 | 106454 | 10927 | 437 | 76 |
|  | full ITS | 316524 | 101794 | 10589 | 413 | 75 |
|  | all records | 370954 | 108567 | 11049 | 441 | 76 |
| rescue sequences | ITS1 | 336786 | 106667 | 10906 | 417 | 75 |
|  | ITS2 | 379015 | 108440 | 11057 | 440 | 77 |
|  | full ITS | 316524 | 101794 | 10589 | 413 | 75 |
|  | all records | 399277 | 111201 | 11218 | 444 | 77 |
| GBIF taxonomy | ITS1 | 335698 | 98200 | 9914 | 393 | 76 |
|  | ITS2 | 377885 | 99644 | 10075 | 415 | 78 |
|  | full ITS | 315505 | 94010 | 9663 | 389 | 76 |
|  | all records | 398078 | 102046 | 10201 | 419 | 78 |
| dereplication | ITS1 | 245596 | 98200 | 9914 | 393 | 76 |
|  | ITS2 | 264245 | 99644 | 10075 | 415 | 78 |
|  | full ITS | 232045 | 94010 | 9663 | 389 | 76 |
|  | all records | 277796 | 102046 | 10201 | 419 | 78 |
| fungal filter | ITS1 | 245526 | 98190 | 9912 | 393 | 76 |
|  | ITS2 | 264159 | 99629 | 10072 | 415 | 78 |
|  | full ITS | 231996 | 94002 | 9661 | 389 | 76 |
|  | all records | 277689 | 102029 | 10198 | 419 | 78 |
| distance based filter & ambiguous nucleotide filter | ITS1 | 239815 | 96320 | 9650 | 343 | 75 |
|  | ITS2 | 258010 | 97720 | 9787 | 360 | 76 |
|  | full ITS | 226407 | 92111 | 9386 | 339 | 74 |
|  | all records | 271418 | 100132 | 9919 | 363 | 77 |

### Cluster threshold impact on identification success

Figure 3 shows how success in species identification using the three ITS databases is impacted by clustering thresholds. Considering only those species with at least two ITS sequences (191,927 ITS1; 219,396 ITS2; 176,714 full ITS) against clusters created from the same data, reduced success in species identification as clustering thresholds were decreased (Figure 3). For example, clustering at 97% identity led to a species assignment for <25% of the queries, roughly half the success for an unclustered (or clustered at 100% identity) database (Figure 3). Considering all three ITS sequence sets, the percentage of identified sequences dropped at least twice as fast as the error rate for species identification for each clustering step (Figure 3). The full ITS region is more sensitive to clustering, as a clustering approach by similarity percentages affects more nucleotides due to its length, disproportionally diminishing the identification gain over ITS1/2 even at the smallest clustering step (Figure 3).

**Figure 1:**
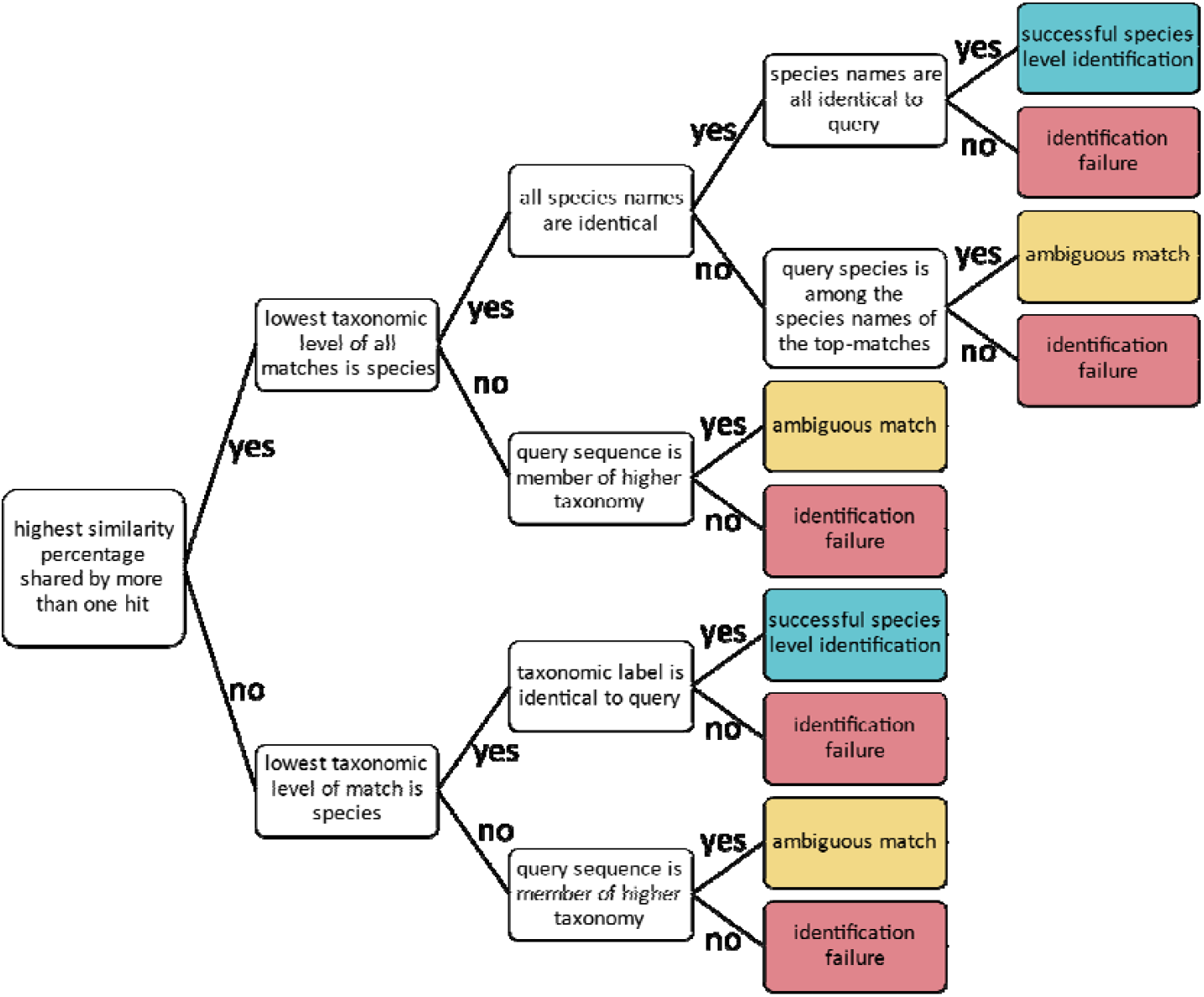
Flowchart of match evaluation logic

**Figure 2:**
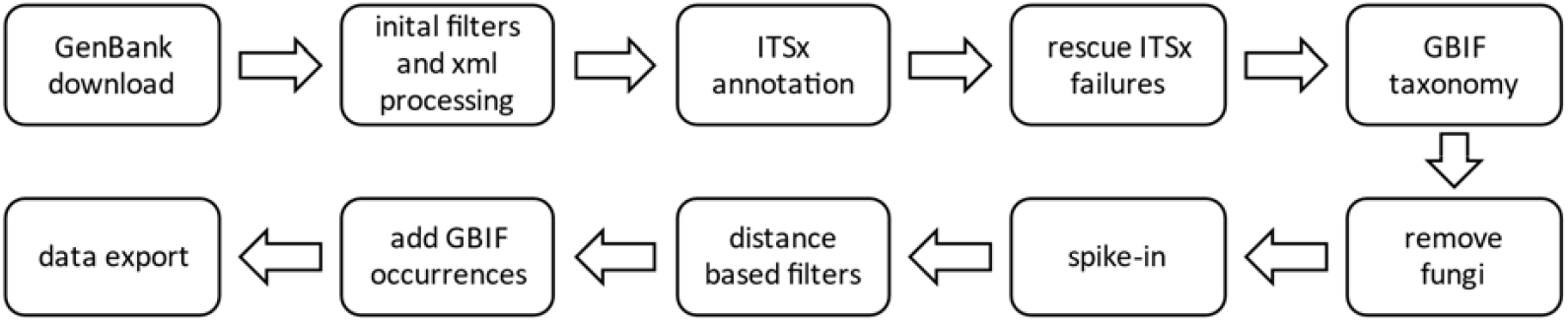
Flowchart of the data curation pipeline used to harvest and validate ITS data from GenBank.

**Figure 3:**
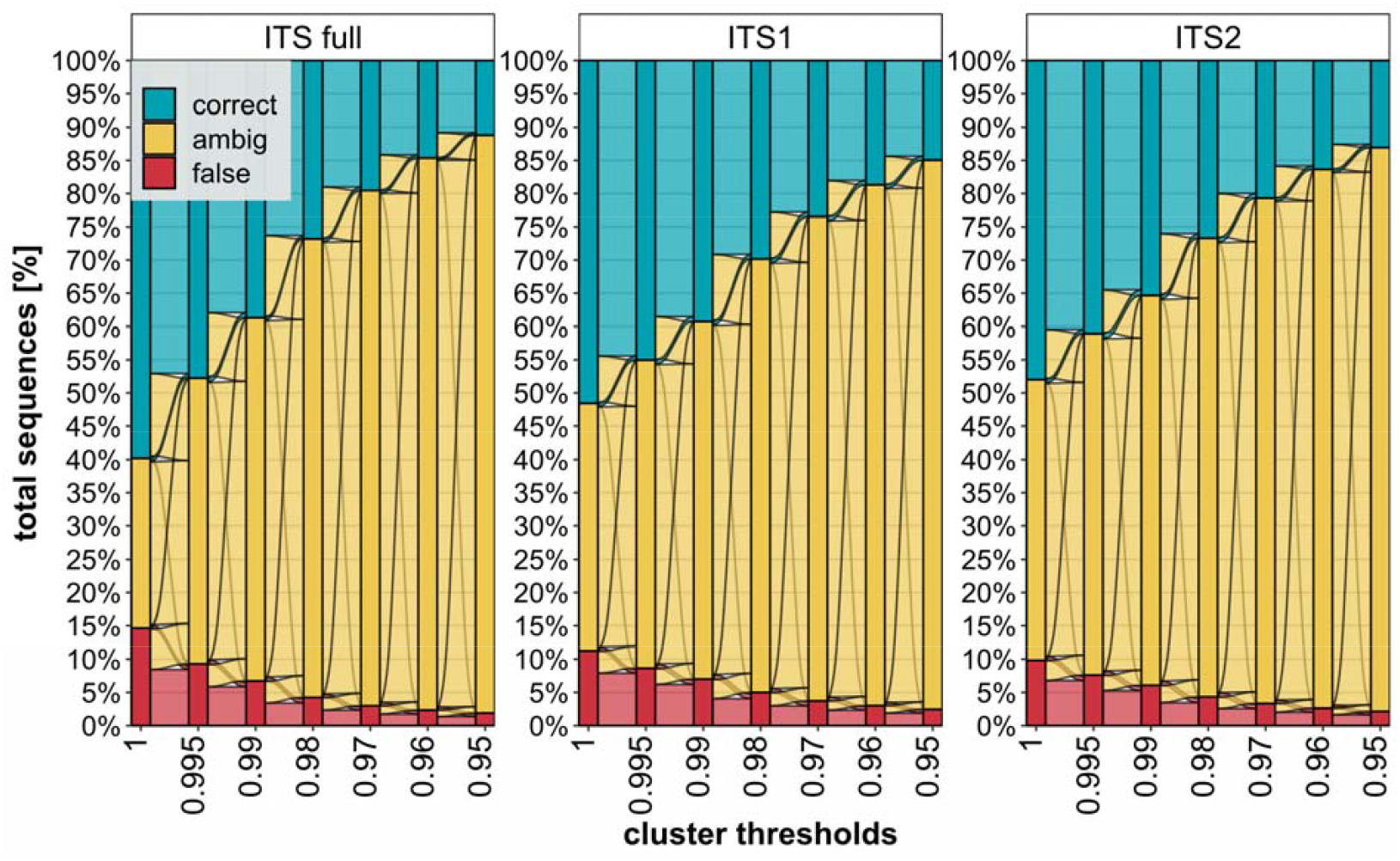
Alluvial plot of identification success at different clustering thresholds. A value of 100% corresponds to 191,927 (ITS1), 219,396 (ITS2), and 176,714 (full ITS) sequences (y-axis) with a minimum of two sequences per species. Cluster thresholds (x-axis) are the minimum similarity value required for two sequences to join a cluster. Colored transitions represent the change from one classification (correct, ambiguous, false) to another between neighboring data points (x-axis).

### Similarity threshold impact on species-level identification success

Figure 4 shows the impact of rejecting reference database matches for varying similarity thresholds using a database with a cluster threshold of 1, representing all exact sequence variants (ESV). The first analysis which is based on species with at least two ITS sequences, shows that only accepting reference database matches with a similarity higher than 0.99 leads to a higher loss of successful identifications than of erroneous identifications (Figure 4A). The use of lower acceptance thresholds (0.98, 0.97, 0.96 and 0.95) has only a minor impact, changing error and identification rates by <1% (Figure 4A). Depending on the ITS region, a rejection threshold of 0.98 removes about 5% of all evaluated matches with the split between removed erroneous and successful identifications being near equal (Figure 4A).

**Figure 4:**
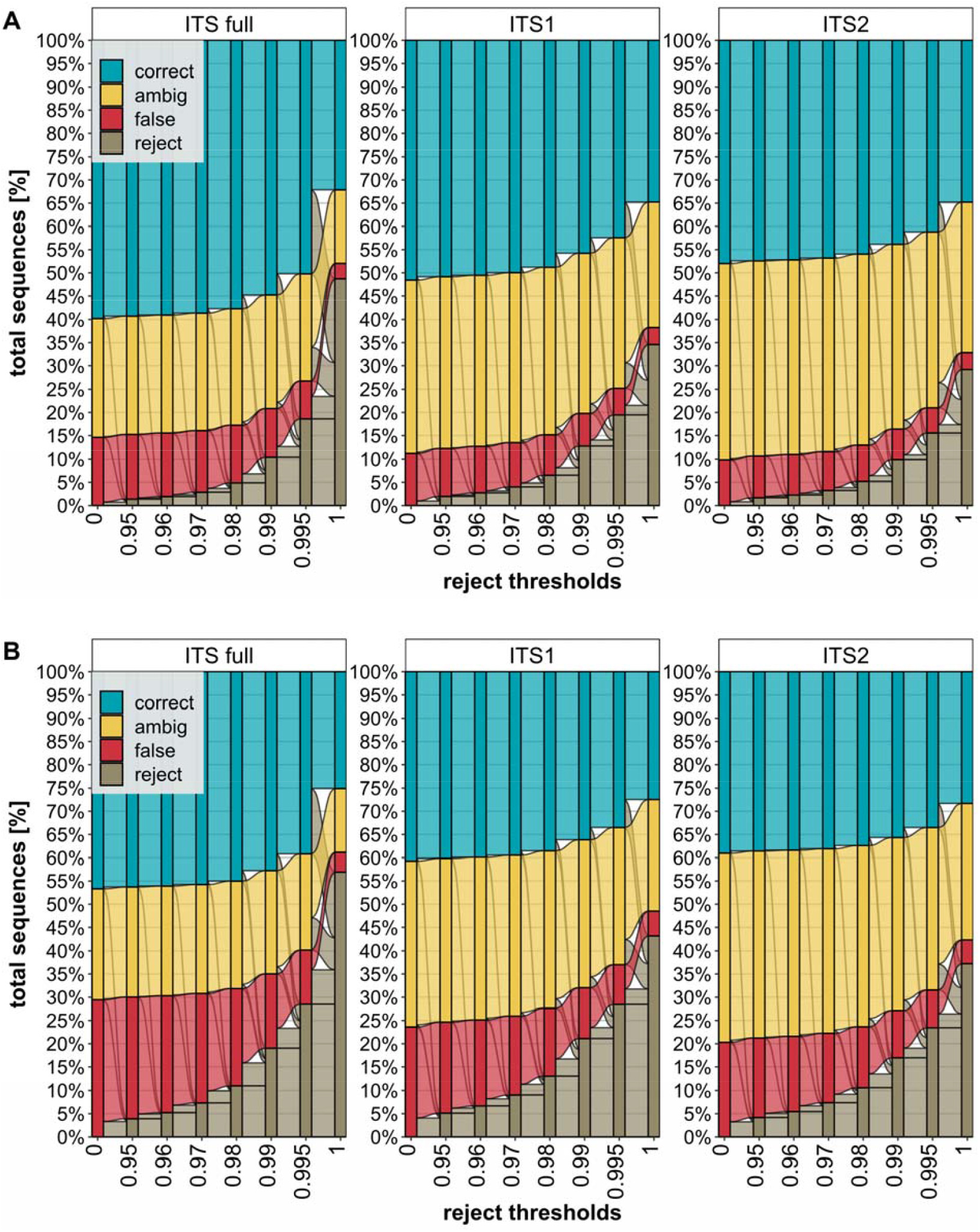
Alluvial plot of identification success at different match distance thresholds. Rejection threshold (x-axis) is the minimum required similarity for a match to be considered. **A:** A value of 100% corresponds to 191,927 (ITS1), 219,396 (ITS2) and 176,714 (full ITS) sequences (y-axis) with a minimum of two sequences per species. **B:** A value of 100% corresponds to 243,015 (ITS1), 269, 675 (ITS2) and 226,420 (full ITS sequences (y-axis), including species represented by one sequence. Colored transitions represent the change from one classification (correct, ambiguous, false) to another between neighboring data points (x-axis).

The second analysis employed an expanded dataset, as it included species with only one sequence (∼54% of total species). Because the removal of self-hits prevents the latter species from receiving a correct match, the error rate is twice as high as in the previous analysis (Figure 4). At all rejection thresholds, except 1, more erroneous identifications are removed than successful identifications (Figure 4B). For full ITS a threshold of 0.995 rejected 28.5% of all matches. Approximately 18% of the excluded matches were erroneous identifications, 7.5% were correct species assignments and 3% were ambiguous assignments. Overall, this decreases the error rate from ∼29.5% to ∼11.5%, comparable to the 12.5% of the previous analysis where species represented by one sequence were excluded at a reject threshold of 0.98 with 5% total rejections (Figure 4).

## Discussion

### Evaluation and choosing of ITS plant reference databases

Multiple workflows have been used to create ITS reference databases for plants, but there is no objective metric to compare their performance. The reference database with the most species will inevitably be the one that has received the least amount of curation. The database(s) with the highest entropy or with the most unique sequences will have employed the most relaxed filtering for pseudogenes (Dubois et al., 2022). The database with the highest sequence count after dereplication will simply be the one which has been downloaded from GenBank most recently. The database with the highest rate of successful identifications has been generated by most aggressive filtering after cross-validation, using a priori taxonomic knowledge, but this approach potentially removes valid sequence variation within a species. Ideally, the database which best represents the intraspecific diversity of any given species should be employed. However, that is unverifiable due to the low number of sequences per species in GenBank. To ensure the quality of this database, we implemented the following five steps to address the plant-specific challenges in the use of ITS for barcoding:

#### (1) Use ITSx for Annotation

Although primarily designed to annotate individual segments of the ITS array (18S, ITS1, 5.8S, ITS2, 26S), it can be used to identify and remove pseudogenes (Buckler et al., 1997; Harpke and Peterson, 2008; Bengtsson-Palme et al., 2013). Substitutions in conserved regions can cause annotation failures due to mismatches with the HMMER profiles used by ITSx (Finn et al., 2011). Additionally, user-defined taxonomic limitations in ITSx’s HMMER profiles can help to exclude sequences from non-target organisms, such as fungi.

#### (2) Adopt strict thresholds for 5.8S

The conserved 5.8S region has been used as a diagnostic tool for decades and is an important tool in post-analytical validation (Jobes and Thien, 1997) both for intraspecific genetic distance and sequence length, optimize filter performance.

#### (3) Employ fixed thresholds for dereplication

instead of keeping all unique sequences this approach does not exacerbate ITS sequence diversity by treating rare pseudogenes and functional ITS variants equal. Especially rare ITS variants from ancient hybridization events, which can occur even across multiple genera, complicate barcoding (Mahelka and Kopecký, 2010).

#### (4) Identify and remove incorrectly annotated plant sequences

adding correctly annotated fungi accounts for non-plant amplification due to low plant-specificity in some ITS primers (Hollingsworth et al., 2011; Kolter and Gemeinholzer, 2021b).

#### (5) Relax intraspecific distance-based filtering

attributes to the fact that ITS sequences from one specimen might present themselves in divergent variants, e.g. forming non-monophyletic species (Xu et al., 2017). Although a more stringent filtering potentially detects more taxonomically incorrectly named sequences, the number of sequences per species, on average, is well below 3, making it difficult to find statistically robust means of excluding sequences (Kolter and Gemeinholzer, 2021a). It also cannot be assumed that the majority of sequences of any given species are correct due to widespread systematic errors across multiple studies, such as in the genus Equisetum (Ibi et al., 2022).

### Incorporating plant occurrence information

This workflow includes occurrence data from GBIF but does not generate regional subsets of the database. Instead, we opt for a post-analysis verification step defined by extracting occurrence information from metadata and checking the plausibility of barcoding results manually.

Although the exclusion of plant species absent from a particular study area can raise the success of assigning sequences to a plant species, it has limitations. On average, just55% of the species in plant lists for European nations possess ITS barcode sequences on GenBank, limiting identification success (Quaresma et al., 2023). Moreover, the completeness of national plant lists is unverifiable, and they often exclude introduced species. For instance, GBIF records fail to include commonly cultivated plants (e.g., doi.org/10.15468/dl.8u8d6h). Contaminating sequences (e.g., *Gossypium, Citrus*), also might be absent from national reference databases, further complicating identification. For these reasons, restricting a barcode reference database, using potentially incomplete national plant lists in combination with incomplete barcode coverage might result in an increased rate of unverifiable false-positive identifications masquerading as a species-level identification increase.

Despite concerns about the reliability of GBIF occurrence data, many criticisms do not apply here (Zizka et al., 2020). Filters based on inaccurate coordinates, duplicate records, incorrect dates, sea/land discrepancies, urban areas, and unknown collection methods are irrelevant for this study, as the data is aggregated at the country level, and the sampling method is not a concern. Including urban areas is necessary for environmental DNA studies. The only problematic records are herbarium records incorrectly attributed to storing institutions instead of their actual occurrence. Disjunct distributions, such as those combining Asia, the UK, and the USA, should be checked for plausibility. Automatic filters are not feasible, as many invasive species are found in EU or North American countries.

In conclusion, we propose a post-analysis alternative to database subsets, enabling researchers to make informed decisions rather than excluding matches based on unverifiable parameters. Unlike static plant lists, this workflow enables automatic occurrence data retrieval during execution of the script.

### Clustering and distance thresholds

Clustering of ITS sequences has been reported multiple times (Banchi et al., 2020; Wirta et al., 2021; Namin et al., 2022). However, studies comprehensively analyzing the impact of such clustering on a larger scale are missing. Our analysis shows that clustering with a similarity threshold smaller than 1 (ESV) reduces identification success disproportionally to the reduction of erroneous identifications and should hence be avoided (Figure 3).

The enforcement of distance thresholds for identification, an approach which rejects matches between a query sequence and records in the database when a defined similarity value falls below a threshold, has also been employed, but without a plant-specific study validating this approach (Parveen et al., 2016; Lucek et al., 2019; Bänsch et al., 2020). Our study reveals that the adoption of an identification threshold, using a global plant database, can reduce the misclassification in species-level plant identifications under certain conditions. If all anticipated query species are present in the reference database, there is no need to set a threshold, but it should not be stricter than 0.98 if adopted (Figure 4A). In a more realistic scenario, using a 0.995 threshold on a reference database missing half of the query species, we rejected ∼28% of matches, achieving an error rate similar to that of a complete database (Figure 4B). Generally, we considered a threshold beneficial if the number of rejected identifications that would have otherwise scored as successful is not higher than the number of rejected identifications which otherwise would have been erroneous.

Application of a universal threshold across Tracheophyta, due to the variation in intra- and interspecific distances of ITS sequences, always introduces errors. The recommendation to use a specific threshold across all Thracheophytes is always a stochastic trade. However, in our case, on average across all plant sequences included in this study, the reclassification of sequences based on our suggested threshold of 0.995, using an incomplete database, results in a correct reclassification in 3 out of 4 cases and introduces a false negative in 1 out of 4 cases. The false negative errors introduced are approximately 25% (7.5% out of 28%). In our use case we could demonstrate that no additional false positive errors do gets introduced by this treatment, as none of the previously successful identifications were transformed into erroneous (red) identifications (Fig. 4). Whether or not this is acceptable has to be evaluated based on the question behind the respective study at hand.

Applying a universal threshold across Tracheophyta introduces errors due to variations in intra- and interspecific distances of ITS sequences (Kolter and Gemeinholzer, 2021a). In our study, reclassification based on a 0.995 threshold, using an incomplete database, correctly reclassifies affected sequences in 3 out of 4 of cases and introduces false negatives in 1 out of 4 of cases (7.5% out of 28%, see Methods). Using this fixed threshold is a stochastic trade-off which on average, improves the identification reliability while not introducing additional false positive errors. While in the case of diverse samples, such as environmental DNA, this stochastic approach most likely will not result in any disproportionally negative impact, we hypothesize that targeted studies, limited to a specific set of taxa, should re-evaluate the threshold proposed here.

## Conclusion

By coupling sequence information with occurrence data and GenBank metadata, the present script allows researchers to validate their identifications using multiple metrics. The unattended script execution provides a dynamic workflow, ensuring up-to-date reference databases. Our analysis showed that clustering of records leads to a substantial reduction in the success of species assignment, but identification thresholds were useful in some situations. Future improvements, such as more sophisticated filtering techniques, depend on closing gaps in reference databases and increasing the average number of sequences per species. Retaining the flanking sequence information (18S & 28S) from GenBank makes this workflow suitable for future long-read barcoding reference database creation (e.g., Nanopore).

## Methods

The R script developed in this study is available on GitHub together with detailed usage notes (https://github.com/Andreas-Bio/ACVPMBD). R package snapshot information, created by the R package renv, is available to increase reproducibility (Ushey and Wickham, 2023). All steps can be run without user intervention (Figure 2). Whenever the script is executed, a pre-set query, which can be modified by the user, determines which GenBank records will be downloaded by the R package rentrez (Winter, 2017). Downloaded sequences without a species-level assignment are discarded while other metadata is retained in tabular format and the sequence is formatted using the R package Biostrings (Pagès and Aboyoun, 2017). Initial filters also remove sequences with illegitimate characters in their species name, sequences with more than 2% ambiguous nucleotides, and sequences shorter than a customizable minimal length.

If present, the 26S, ITS1, 5.8S, ITS2, and 18S regions are annotated, as detected by ITSx (Bengtsson-Palme et al., 2013). Chimeric sequences detected by ITSx are excluded and sequences that fail to be annotated for either ITS1 or ITS2 are also removed. ITSx needs a small segment of the gene regions adjacent to ITS1/2 to function properly. However, some data uploaded to GenBank only includes the core ITS1/2 regions as the flanking areas have been trimmed. To address this, a recovery effort is made by comparing successfully annotated ITS1/2 regions with other sequences. Unannotated sequences with a global (including terminal gaps) similarity of >85% to the annotated sequences are retained while the others are discarded. Sequence information and metadata are combined into a tab delimited database file. Each dataset record is uniquely identified by its GenBank accession number, and it can have up to six associated sequences (ITS1, ITS2, full ITS, 26S, 18S, 5.8S).

The taxonomy assigned to the sequence records extracted from NCBI is compared to the GBIF backbone taxonomy (doi.org/10.15468/39omei). In detail, a query, encompassing taxonomic levels from species (with author’s epithet) to kingdom as per NCBI, is matched against the GBIF taxonomy backbone using the R package rgbif (Chamberlain and Boettiger, 2017). In cases of discordances, the GBIF taxonomy has priority and the taxonomic assigned by NCBI is replaced, but still retained as metadata.

The number of sequences for each species is then determined by a dereplication step with a default of 10(Kolter and Gemeinholzer, 2021a). Priority for retention is first given to full length ITS sequences (as determined by ITSx) and then to different GenBank accession identifiers (first 3 characters) with the aim of retaining as much genetic variation as possible, as different projects are usually assigned different alphanumerical accession identifiers.

Sequences are then examined to remove those reflecting contamination. To aid the recognition of contaminants, a library of outgroup sequences was established. This was accomplished by extracting one sequence from each fungal genus in the UNITE database (version 9.0) and adding a manually curated subset of sequences for Bryophyta and Algae from GenBank (Abarenkov et al., 2023). Based on a dynamic distance threshold, designed based on data from a previous study (Kolter and Gemeinholzer, 2021a), sequences matching any outgroup sequence were removed from the ITS databases. Outgroup sequences were secondarily incorporated into the database to act as a sink for sequences from non-target taxonomic groups recovered by plant barcoding or metabarcoding studies.

Multiple distance-based filtering steps were incorporated to recognize probable labelling or identification errors. First, using a leave-one-out-cross-validation (LOOCV), the family of the top hit for each sequence was compared with that assigned to the query. When a mismatch was detected, both sequences (query, match), were again queried against the remaining records in the database. If the mismatch persists, the query sequence is removed. This step also leads to the removal of families represented by a single sequence. All following distance-based filtering steps were also applied to the 5.8S region with tightened thresholds, due to its conserved nature. A second LOOCV filtering step removes query sequences from the database if the species with the top hit does not match the query species and the median distance of the hits to the query species is greater than a set threshold (default: 10%, 5.8S: 5%). In the third filtering step, any sequence not matching any other sequence within a defined threshold (default: 30%, 5.8S: 10%) is removed from the database. Finally, every region (separately: ITS1/2, full ITS, 5.8S) with more than a set threshold of ambiguous nucleotides (default: 2.5%, 5.8S: 1%) is removed from the database.

Distributional information for each species is added to the database based on GBIF occurrence data. The countries along their respective occurrence count, if above three, are reported in alphabetical order and grouped by the United Nations region definition.

The script outputs the whole database in a tab delimited format, as a reduced database without sequence information, and for any specified ITS region as a fasta file. An ITS sequence length plot, summarized by family, and an interactive taxonomic summary plot are automatically generated (Ondov et al., 2011). In addition to a repository on Zenodo (10.5281/zenodo.10257823), which includes all temporary files, the database has also been uploaded to BOLD (project code: PREF), which makes the data available through the publicly accessible identification engine.

### Cluster threshold impact on identification success

Clusters were generated from three sequence files (ITS1, ITS2, full ITS) using the script described earlier for seven similarity thresholds (1, 0.995, 0.99, 0.98, 0.97, 0.96, 0.95). Viewed from the perspective of sequence matching, a clustering threshold of 1 generates an identical result to an unclustered database if multiple top matches are evaluated. The clusters were given new labels based on the lowest taxonomic lineage shared among members of each cluster.

Sequences from species represented by at least two sequences (45,302 ITS1; 47,677 ITS2; 45,302 full ITS) were matched against clusters of the respective ITS region at each threshold level using VSEARCH (Rognes et al., 2016). Self-hits were removed by eliminating the query sequence from the matching clusters followed by updating the taxonomic label of the respective cluster. The match or matches with the highest percent similarity, excluding terminal gaps, were defined as top-hit(s) and were evaluated for a taxonomic match. If the top-hit(s) all belonged to one species and were identical to the query sequence, the match was considered a successful species-level identification. If the top-hit(s) included multiple species, one identical to the query, or if the top hit(s) had a higher taxonomic level than species (i.e., cluster only labeled to genus because it contained multiple species), but if the query species was a member of the respective higher taxonomic level, the match was considered ambiguous. If the query species was not among the species of the top-hit(s) or the query species was not a member of the higher taxonomy of the matching cluster(s), the match was classed as an unsuccessful species-level identification (Figure 1). A total of 4,116,117 evaluations, across all combinations, were combined in an alluvial plot by the R packages ggplot2 and ggalluvial (Brunson, 2020; Wickham, 2016).

### Threshold based rejection of database matches

We re-analyzed matches between ITS sequences and clusters generated with a similarity threshold of 1 (ESV) generated earlier. In contrast to the evaluation discussed earlier, two query sets were used: one with species having at least two sequences, and another including all species including those represented by one sequence. As self-hits were removed for both query sets, this increased the incidence of incorrect or ambiguous match assignments if the query species was eliminated by this step, simulating an incomplete database. Top-hits with a distance (nucleotide similarity percentage without terminal gaps) below seven similarity thresholds (1, 0.995, 0.99, 0.98, 0.97, 0.96, 0.95) were categorized as rejected. These steps were repeated for each ITS region, for each threshold and for both query sets. All other evaluation and visualization steps were identical (see above).

## Data Accessibility Statement & Benefit-Sharing Statement

Scripts and data used to create the results presented in this publication are available from: https://github.com/Andreas-Bio/ACVPMBD. All output files for all filtering steps and the final reference data files can be downloaded at DOI: 10.5281/zenodo.10257823. Benefits Generated: Benefits from this research accrue from the sharing of our data and results on public databases as described above.

---

**Supplementary Table 1:**
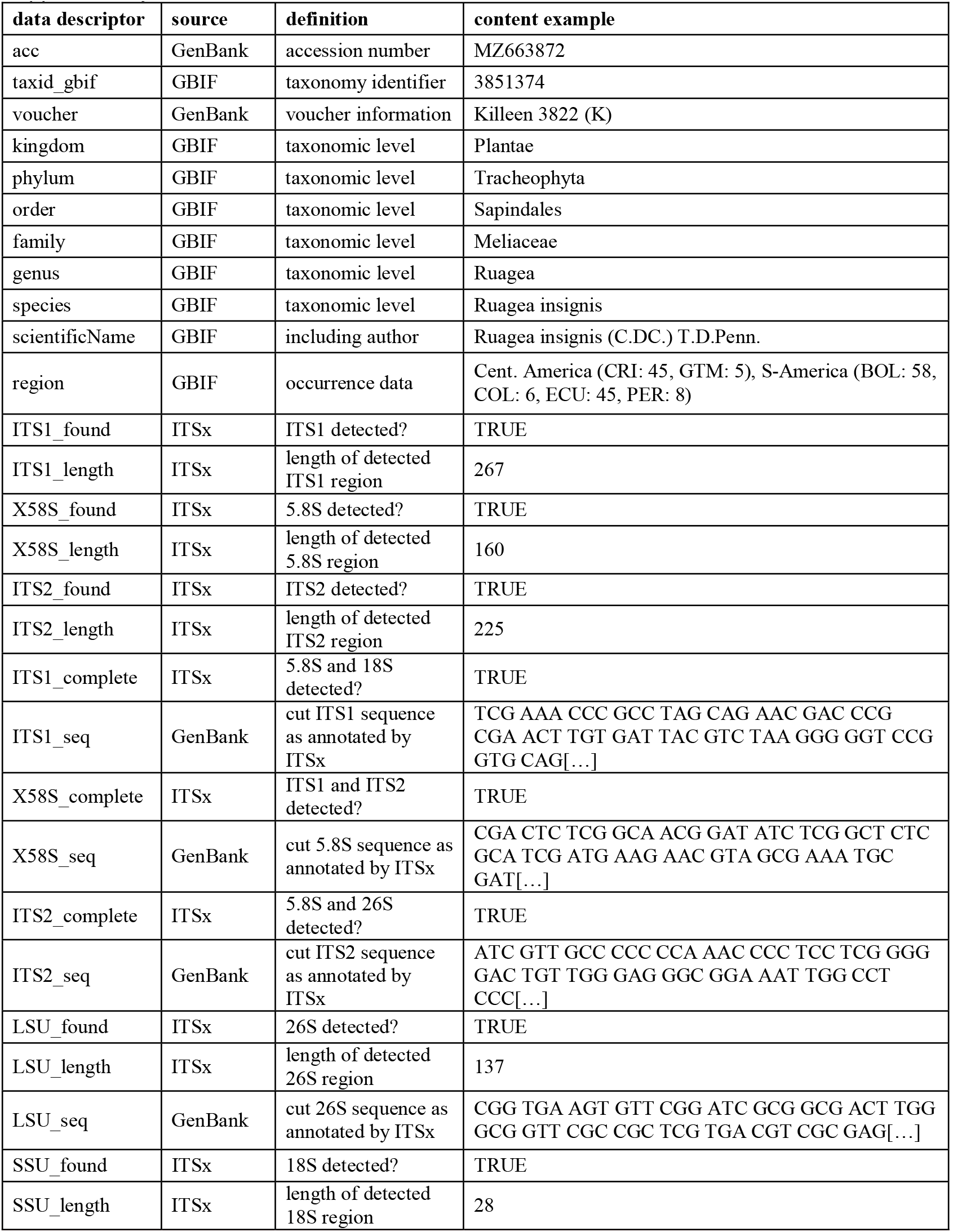
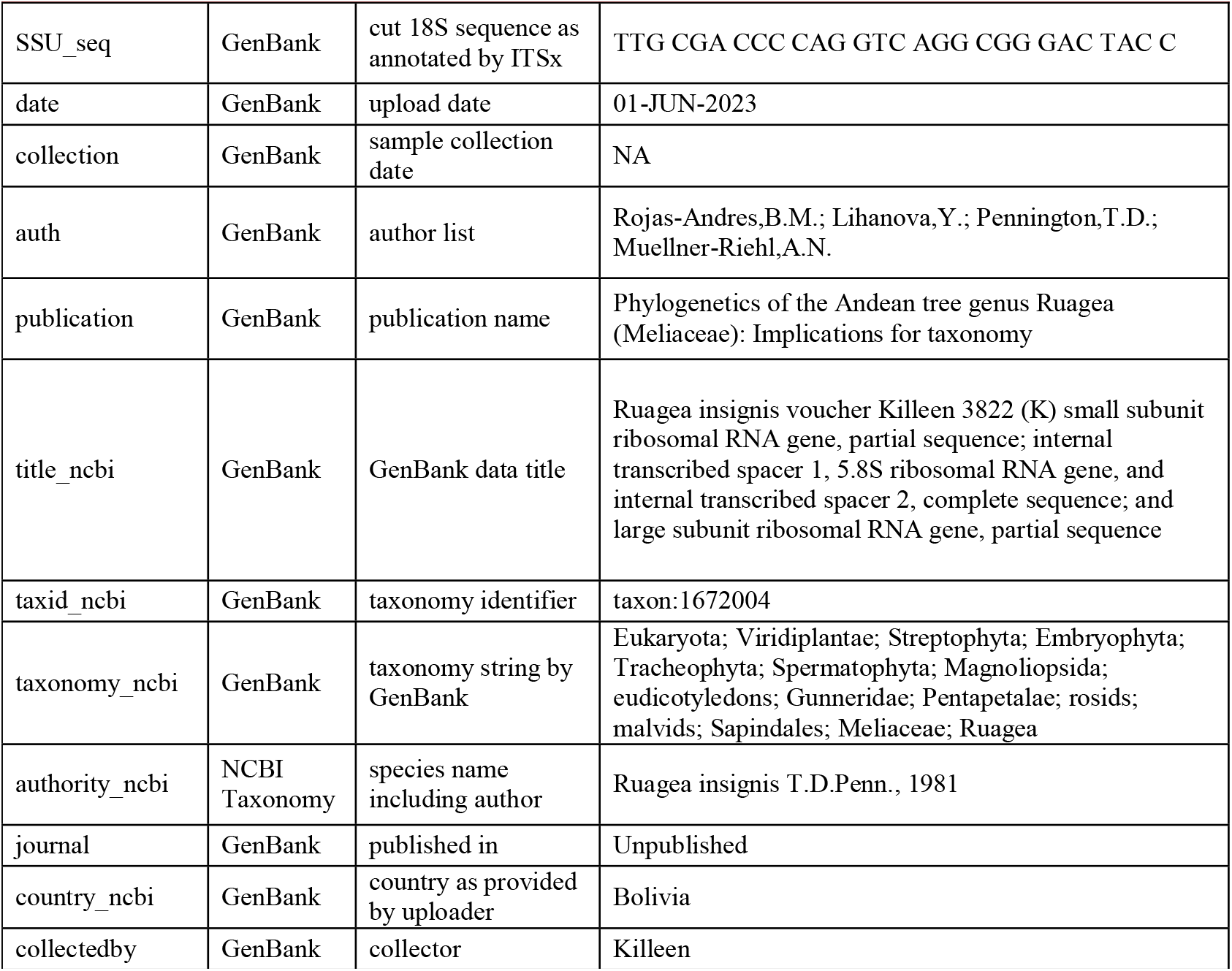
Data records stored in the reference database in a tabular format.

